# A method for focused ultrasound (FUS) neuromodulation with simultaneous electroencephalogram recordings in awake, head-fixed mice with temporal lobe epilepsy

**DOI:** 10.1101/2024.03.20.585988

**Authors:** Carena Cornelssen, Braden Brown, Eli Finlinson, Skyler Blair, Sharayu Senthilkumar, Jan Kubanek, Karen S. Wilcox

**Affiliations:** Department of Biomedical Engineering, University of Utah, Salt Lake City, Utah, USA; Department of Pharmacology and Toxicology, University of Utah, Salt Lake City, Utah, USA; Department of Electrical & Computer Engineering, University of Utah, Salt Lake City, Utah, USA; Interdepartmental Program in Neuroscience, University of Utah, Salt Lake City, Utah, USA

**Keywords:** focused ultrasound, epilesy, neuromodulation, awake, head-fixed mice, low-intensity ultrasound

## Abstract

Transcranial focused ultrasound (FUS) may be a promising neuromodulation technology for treating people with epilepsy whose seizures are drug resistant. Prior studies have shown seizure suppression in animal studies using FUS. However, most of these studies were performed in evoked seizure models and not in animal models of epilepsy. Evoked seizure models do not exhibit the pathophysiology of epilepsy and do not exhibit spontaneous recurrent seizures, which define epilepsy. For translation to humans, there is a critical need to determine the specific FUS stimulation parameters that reduce spontaneous recurrent seizures in a chronic disease model of epilepsy. To achieve this goal, we developed and optimized an approach to determine the effects of ultrasonic stimulation on metrics of seizure-like events (SLEs) in awake, head-fixed mice within the intrahippocampal kainate (IHK) mouse model of temporal lobe epilepsy (TLE). A proof-of-principle study demonstrated that two target (bilateral and contralateral to the kainic acid injection site) stimulation conditions and two FUS parameter sets (low and high pressure) could be combined with the ability to simultaneously record hippocampal electroencephalograms. We also provide a method for analysis of the effects of FUS stimulation on the metrics of SLEs (interevent duration, SLE duration, and spike frequency).

## Introduction

Epilepsy is a common neurological disorder characterized by recurrent spontaneous seizures and affects an estimated 65 million people worldwide (Hesdorffer et al., 2011). The most common form of refractory (drug-resistant) epilepsy is temporal lobe epilepsy (TLE) (Engel et al., 2012). Neuromodulation approaches to treat seizures accompanying epilepsy often requires targeting sub-cortical brain targets (Mahoney et al., 2020). Deep brain stimulation is spatially specific and can reach sub-cortical brain targets but is an invasive therapy (Mahoney et al., 2020). Modern treatments like transcranial magnetic stimulation are noninvasive, but do not reach sub-cortical brain targets and there is limited evidence that this approach can reduce the frequency of seizures (Mahoney et al., 2020; Walton et al., 2021). Transcranial low-intensity focused ultrasound (FUS) is a neuromodulation technology that is spatially specific, noninvasive, and can target sub-cortical structures, which are often targets for treatments in TLE (Mahoney et al., 2020). Therefore, FUS may be a promising treatment for refractory epilepsy (Cornelssen et al., 2023).

FUS has been shown to suppress seizures in rodent and nonhuman primate studies (Min et al., 2011; Hakimova et al., 2015; Li et al., 2019b, 2019a; Chen et al., 2020; Lin et al., 2020; Zou et al., 2020; Murphy et al., 2022). However, most of these papers did not investigate FUS in a chronic model of epilepsy, using evoked seizure models instead, which made the results difficult to translate to the disorder state (Min et al., 2011; Hakimova et al., 2015; Li et al., 2019b, 2019a; Chen et al., 2020; Lin et al., 2020; Zou et al., 2020). Additionally, many studies used anesthetized animals, making it difficult to translate the findings to the awake state in a model of epilepsy (Min et al., 2011; Li et al., 2019b, 2019a; Chen et al., 2020; Lin et al., 2020; Zou et al., 2020). Therefore, there is a critical need to determine the specific FUS stimulation parameters that can modify seizures in the chronic state of TLE. There is also a need to develop a method to study FUS stimulation parameters in awake, head-behaving animals prior to human translation.

Our goal was to establish a FUS technique in awake, head-fixed mice while simultaneously recording electroencephalograms (EEGs). As proof of principle, we tested our approach in mice to identify the effects of two FUS stimulation parameters on metrics of SLEs in the intrahippocampal kainate (IHK) mouse model, a well-validated preclinical chronic model of TLE (Duveau et al., 2016; Twele et al., 2017). This method can also be utilized in other mouse models of neurological disorders, such as depression and anxiety.

## Methods and Materials

### Animal surgery

The University of Utah Institutional Animal Care and Use Committee approved all animal procedures and ARRIVE Guidelines were followed. Seventeen male C57BL/6 mice (4-5 weeks old; Jackson Labs) underwent stereotactic brain surgery under isoflurane anesthesia (1-5%) to induce status epilepticus (SE; Figure 1). A half-millimeter (mm) diameter-sized craniotomy was created by drilling a bur hole using a hand drill (Dremel 2050-15 Stylo+) with a spherical bur (19007-05, Fine Science Tools). Next, either 20 millimolar (mM) of kainic acid (KA; Tocris Bioscience) or sterile saline (0.9%, Medline Sterile Saline) was injected at a rate of 50 nanoliters (nL)/min for two minutes for a total volume of 100 nL into the dorsal hippocampus of the mice using a micro syringe pump controller (Micro4™, World Precision Instruments). Borosilicate glass (20443936, World Precision Instruments) micropipette electrodes were used for injection of KA or saline and were pulled using an electrode pipette puller (P-97 Flaming/Brown Micropipette Puller, Sutter Instrument) to a taper of 15-20 micrometer (μm) thickness. Mice were injected with either KA (n=8) or sterile saline (n=9) within 0.2 mm^3^ of the following coordinates: anterior-posterior (AP): −2.00 mm, medial-lateral (ML): 1.25 mm, dorsal-ventral (DV): 2.00 mm based on the standard mouse atlas with their respective electrode implant coordinates (Paxinos and Franklin, 2001).

**Figure 1:**
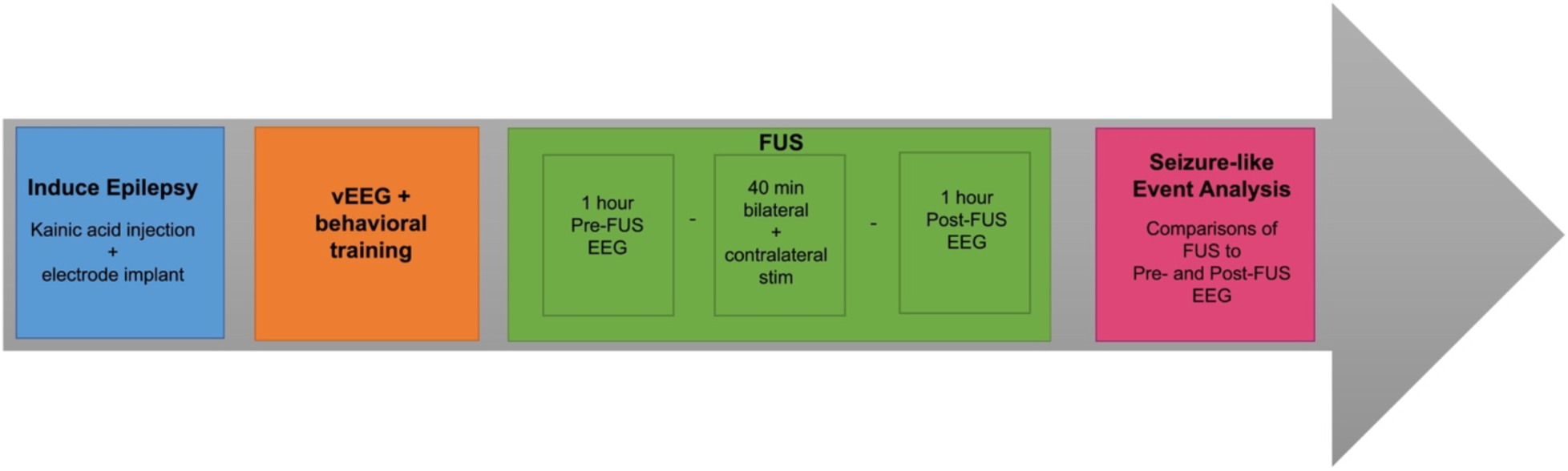
Timeline of the approach. Epilepsy is induced in mice in the blue block with infusion of kainic acid into the hippocampus during weeks 4-5. In the orange block, behavioral training of the mice occurs after recovery from surgery during weeks 8-20. FUS administration (green block) commenced 18 weeks after surgery for the IHK mice (n=2) in the green block. The mice underwent a one-hour Pre- and Post-FUS baseline with 40-minute FUS stimulation periods in the FUS block. The stimulation period included bilateral and contralateral stimulation target conditions. The order of the stimulation target conditions was randomized on the day of FUS administration. The FUS parameter set was randomized each day of the FUS administration and both target conditions were run in a single day. Sessions with different ultrasound parameter sets were run a minimum of three days apart. Data analysis (pink block) commenced at the conclusion of FUS administration.

Following the KA or saline injection, a platinum/iridium recording electrode (8IMS3339BXXE, P1 Technologies), safe for FUS stimulation, was implanted approximately 0.5 mm posterior to the injection site (Figure 2A and 2B) and sealed with super glue (454, Loctite®). The coordinates for electrode placement for all mice (n=17) were within 0.40 mm of the following coordinates: AP: −2.66 mm, ML: 1.75 mm, DV: 1.80 mm. Dental cement (TRIM & TRIM II Liquid, Keystone Industries) was used to seal the implantation sites and the skull and formed a cap on the skull. A custom 3D-printed headplate (acrylonitrile butadiene styrene (ABS) transparent blue 1.75 mm, hatchbox3d) was attached on the dental cement on each mouse’s skull over the occipital and parietal lobes (Figure 3A). This headplate allowed for head fixation for FUS stimulation and behavioral training on a custom-made 3D-printed treadmill (ABS transparent blue and black 1.75 mm, hatchbox3d). Mice were intraperitoneal (i.p.) injected with buprenorphine (0.05-0.10 mg/kg) every 6-12 hours for 72 hours post-surgery for pain during recovery from surgery. As needed, sterile ringer’s solution (Electron Microscopy Sciences) was injected for 72 hours post-surgery to treat dehydration.

**Figure 2:**
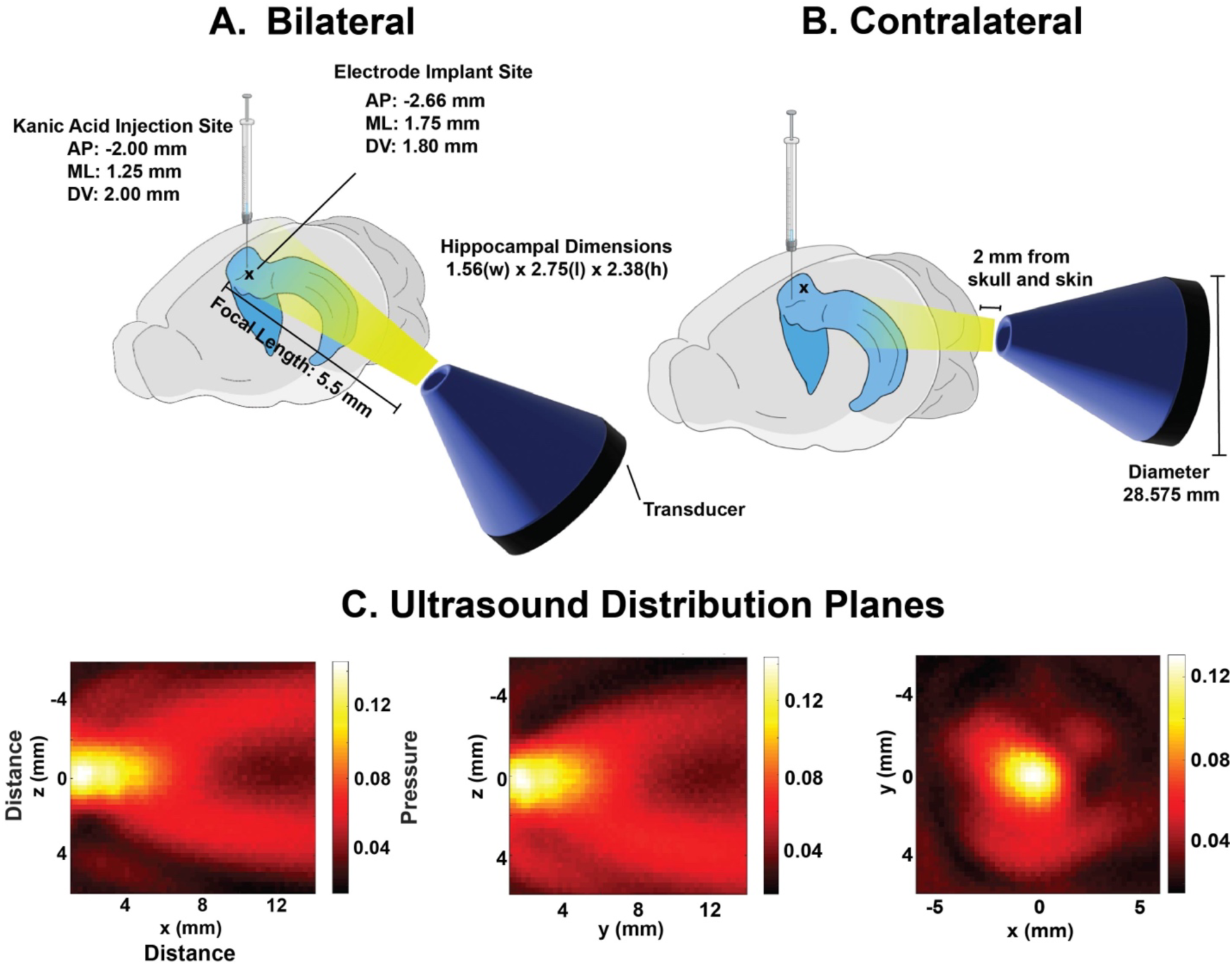
Targets of brain stimulation are shown in ultrasound distribution planes. The projected path of the FUS pressure field shown is shown in yellow in relation to a mouse brain during bilateral (A) and contralateral (B) target stimulation. The target stimulation conditions were named for the FUS pressure focal areas relative to the KA injection site. Bilateral stimulation results in both hippocampi being stimulated whereas contralateral stimulation results in only the hippocampus contralateral to the injection site being stimulated from the transducer placed at the rear of the mouse’s head. (C) Shows the characterization of the ultrasound pressure planes. The ultrasonic pressure field was measured to be 5.5 mm x 4 mm x 3.5 mm (full-width at half maximum measured in a water bath). Stimulation targets were estimated using the measurements of the ultrasonic pressure fields in relation to the standard mouse brain atlas. Additionally, shown in blue, is the position of the collimator tip in relation to the mouse’s head.

**Figure 3:**
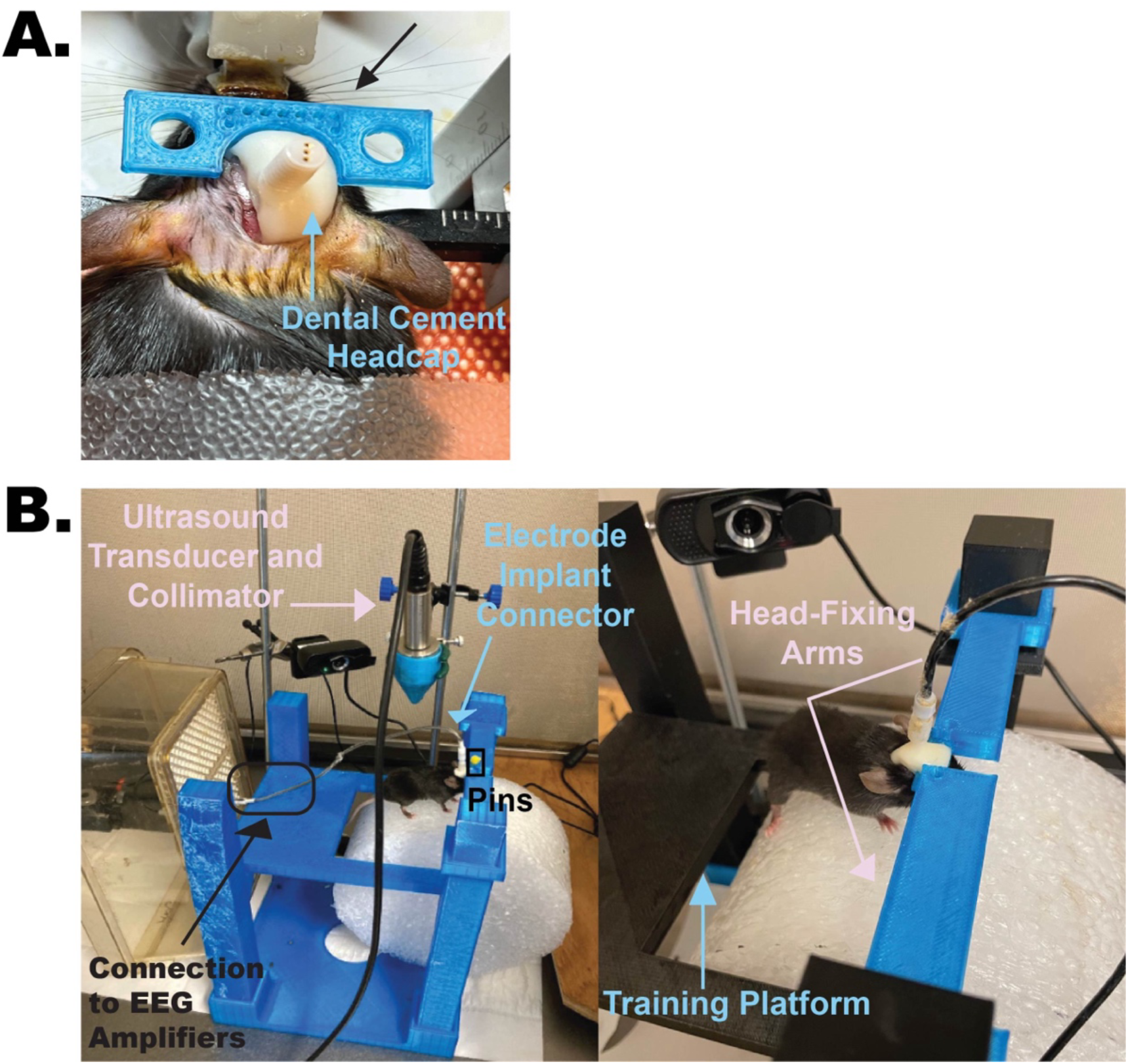
3D-printed treadmill and head-fixing components. (A) The image depicts a mouse during the surgery as the 3D-printed headplate is being added to the mouse’s headcap. The anesthetized mouse is fixed by a stereotactic frame which can be seen on either side of the mouse’s head. The black arrow is pointing to the blue 3D-printed headplate located towards the front of the mouse’s head. The teel arrow is pointing to the dental cement and electrode comprising the headcap. Small holes can be seen in the center of the headplate which were designed to allow the dental cement, yet to be added, to have more surface area to bond to. (B) The custom-made components required for head-fixing an awake mouse during FUS administration while simultaneously recording EEG are depicted.

### EEG and video (vEEG) recordings

Directly following surgery, to confirm the induction or absence of SE following infusion of KA, mice were continuously monitored via EEG recording overnight for 12 hours. Seizure activity was assessed by observing frequent spiking activity and electrographic seizures. After surgery, the mice were provided with a minimum of three weeks of rest and recovery from surgery before undergoing further continuous video and EEG (vEEG) recording. vEEG signals were continuously recorded 24 hours a day, seven days a week (24/7). Electrodes were connected to a commutator (8BSL3CXC0MMT, P1 Technologies) at the top of the cage. This commutator was linked to an EEG amplifier (EG100C, BIOPAC Systems, Inc) via a cable (335441, P1 Technologies). The vEEG amplifier was configured with the following settings: gain set to 1000, a 0.05 Hz filter activated, and a low-pass noise notch (LPN) filter with a notch frequency of 60 Hz to remove noise and a low pass cutoff frequency of 35 Hz. The “norm” option was selected to allow for the visualization of frequency bands passed through the LPN. A custom software was used to record and save the signals at a sampling rate of 500 Hz and record up to 16 channels of vEEG simultaneously (Thomson and White, 2014). Mice in the IHK model not exhibiting spiking activity 12 hours post-surgery, or experiencing complications from surgery or tethering, were not included in the study.

During FUS delivery, vEEG was recorded one hour before the administration of FUS, during the administration of the FUS stimulation for 40 minutes, and one hour after the stimulation. The amplifier and its settings remained consistent with those used during the 24/7 monitoring, except that the sampling rate was altered to 5000 Hz, and two channels were recorded. An EEG recording software (AcqKnowledge 5.0.6 Software, BIOPAC Systems, Inc) package with settings configured to enable two-channel recording facilitated the monitoring of both the EEG and the FUS pulse signals. Additionally, the utilization of a connected camera (XPCAM webcam model N5, PG1600001-04986 HD Webcam 1080P with microphone) was used to capture video footage of the mice during FUS stimulation.

### Design of treadmill and head-fixing plate

A low-cost solution to head-fix the mice was needed during FUS administration. To meet this need, we designed a three-dimensional (3D) printed treadmill with a head-fixing attachment. The treadmill and head-fixing components were custom-made in-house using computer-aided design (CAD) software (Student Fusion 360, Autodesk) and were produced using Fused Deposition Modeling (FDM) and Stereolithography (SLA) 3D printers (Pro2 Plus, Raise3D). The design is available upon request.

### Behavioral training

The behavioral training study included six out of the nine control mice who received saline injections. Based on the cleanest EEG signals, these six mice commenced behavioral training three weeks post-surgery (Guo et al., 2014). IHK mice were enrolled in the training sessions based on whether they exhibited spiking activity. The IHK mice (n=6) in this study underwent behavioral training at least eight weeks post-surgery. Behavioral training was designed to acclimate the mice to walking, running, and being head-fixed to our custom 3D-printed treadmill. The training sessions were conducted within a specially designed area situated inside a Faraday cage within a closed room. Mice were only trained between the hours of 6 AM to 6 PM (Guo et al., 2014). Temperature and humidity in the room were consistent with the housing room. The mice were acclimated to the room at least 30 minutes before training.

While we developed our behavioral training stages, the training process was adapted from Gou et al., 2014 (Guo et al., 2014). During training, we used a sugar reward of 10% sucrose (Sigma-Aldrich) in a deionized water solution in increments of 15-30 seconds (s) to reward the mouse in each stage of the training. On average, mice received 0.5-1 mL of sucrose solution per session. Consistent with results described by Gou et al., 2014, on average mice would have 4-5 sessions a day of stages 1-2, while having 1-2 sessions of stage 3 per day. The behavioral training results were not always linear, as back-tracking occurred in response to the mouse’s behavior. If, at any stage, the mouse exhibited signs of fear or stress, the handler reverted to a previous training stage where the mouse exhibited comfortable behavior, characterized by mastery of the behavioral stage without fear, stress, or struggle. Additionally, the handler increased sucrose reward frequency by decreasing the time between rewards by 5-10 s, providing additional time for the mouse to adapt and feel at ease.

### FUS setup and specifications

A 500-kHz single-element spherical-focused transducer (18-0018-P, Olympus) was employed for all trials. The ultrasound pressure field (Figure 2C) measured 5.5 mm in length, 4 mm in width, and 3 mm in focal height (full-width at half maximum measured in a water bath) when measured through a nine mm diameter-opening of a custom 3D-printed collimator coupled to a C57BL/6 male mouse skull. Measurements were completed using a hydrophone (HGL-0200, ONDA) and hydrophone programmable positioning system (Aims III, ONDA).

The ultrasound transducer was connected to an (55-dB, 258 kHz-30 MHz power) amplifier (A150, Electronics & Innovation) via a 50 Ohm coaxial cable. The amplifier was then connected to a function generator (33500B Series Waveform Generator, Keysight) with two channels. The function generator’s first channel was programmed to output the desired waveforms for the required FUS parameter sets using a MATLAB script (2021a, Mathworks). The function generator’s second channel was linked to an amplifier (EG100C, BIOPAC Systems, Inc), which enabled the synchronous recording of the FUS pulse waveform and the mouse’s vEEG signals.

The FUS stimulation parameter sets used are displayed in Table 1. The low-pressure parameter set used a carrier frequency of 500 kHz, amplitude or pressure of 0.2 MPa, a pulse repetition frequency (PRF) of 500 Hz, a duty cycle (DC) of 50%, a pulse train interval of 0.2 s, an interpulse train interval of 1.8 s, a Mechanical Index (MI) of 0.28, a spatial peak pulse average intensity (I_SPTA_) of 1.33 mW/ cm^2^, and a spatial peak temporal average intensity (I_SPTA_) of 66.67 mW/cm^2^ (Table 1, Low). The high-pressure parameter set used a carrier frequency of 500 kHz, amplitude or pressure of 1.0 MPa, a PRF of 500 Hz, a DC of 10%, a pulse train interval of 0.2 s, an interpulse train interval of 1.8 s, an MI of 1.41, an I_SPTA_ of 3.33 mW/ cm^2^, and an I_SPTA_ of 333.33 mW/cm^2^ (Table 1, High).

**Table 1:** FUS parameter sets used in the FUS administration. The two FUS parameter sets used in the sessions are displayed here, with their carrier frequency (kHz), pressure (MPa), PRF (Hz), DC (%), pulse train interval (s), interpulse train interval (s), MI, I_SPPA_, and I_SPTA_. The parameter sets were designed to examine the impact of high and low intensity FUS stimulation on the suppression of epileptic activity.

| Parameter set | Carrier Frequency (kHz) | Pressure (MPa) | PRF (Hz) | DC (%) | Pulse Train interval (s) | Inter Pulse Train Interval (s) | MI | Isppa W/cm<sup>2</sup> | Ispta mW/cm<sup>2</sup> |
| --- | --- | --- | --- | --- | --- | --- | --- | --- | --- |
| Low | 500 | 0.2 | 500 | 50 | 0.2 | 1.8 | 0.28 | 1.33 | 66.67 |
| High | 500 | 1.0 | 500 | 10 | 0.2 | 1.8 | 1.41 | 3.33 | 333.33 |

### Ultrasound targeting

A 3D-printed collimator made of clear resin (Formlabs) with a 9-mm opening to focus the ultrasound was filled with 4% agar (Kate Naturals). The ultrasound transducer was placed in the 3D-printed collimator and attached with a 3D-printed holder to a micromanipulator (Mitutoyo) to facilitate positioning.

Two stimulation target conditions were used: bilateral and contralateral, each named for the position of the FUS target relative to the injection site in the dorsal hippocampus. The ultrasound transducer was placed at a 5-degree angle from the horizontal for bilateral stimulation condition. The tip of the focusing collimator was then aligned and coupled to the mouse’s head using degassed ultrasound gel (100, Aquasonics®). Prior to use, the gel was further degassed using customized 10 mL syringes (Becton, Dickinson, and Company) with a syringe filter top centrifuged (5702 R, Eppendorf) for five minutes at 2500 rpm. The coupling point was positioned 3 mm from the top of the skull and in line with the electrode. The transducer was positioned using the following: 1) the stereotactic coordinates of the electrode and the injection site (Figure 2A), 2) a standard mouse brain atlas, and 3) the measurement of the ultrasonic pressure field (Figure 2C) (Paxinos and Franklin, 2001). The standard mouse brain atlas was used as a map and marked with the injection site’s stereotactic coordinates. The pressure field, with correct proportions to the atlas legend, was also overlayed on the atlas at different points and angles to ensure the field would reach the injection site coordinates. Based on these measurements, the contact point was determined to be 3 mm from the top of the skull and in line with the injection and electrode.

The transducer was positioned behind the skull near the cerebellum using the micromanipulator for contralateral stimulation. The tip of the collimator was coupled directly behind the mouse’s left ear to avoid stimulation at the injection site (Figure 2B) and the same technique described above was used to ensure targeting of the contralateral hippocampi.

### FUS administration session design

FUS administration commenced 18 weeks post-surgery. Enrollment criteria of a minimum of 10 High-Voltage Sharp-Wave (HVSWs) discharges per hour was used. Two mice from the IHK group met this criterion for inclusion.

Each FUS administration day consisted of two sessions, each with three conditions: 1-hour pre-baseline vEEG, a 40-minute stimulation session with vEEG, and a one-hour post-baseline vEEG. The 40-minute stimulation session consisted of either contralateral or bilateral target stimulation conditions and one of the two FUS parameter sets (low or high) chosen randomly to minimize order effects using an online random number generator. Figure 2 and Table 1 summarize the target conditions and FUS parameter sets given to the mice. Each mouse underwent different FUS parameter stimulation sets every three days. The mice were acclimated to the room for a minimum of 30 minutes before the FUS administration day to become familiar with the environment. Following the session, data analysis occurred offline. This process was repeated on the FUS administration day for the second session but alternating to the second target condition for the mouse using the same FUS parameter set.

### Seizure detection

To ensure the proper selection of mice for the proof-of-concept, clear criteria for SLEs were established. HVSWs are SLEs that are commonly observed in the IHK mouse model (Twele et al., 2016a, 2016b, 2017; Zeidler et al., 2018). HVSWs were defined as a series of high amplitude spikes with a spike frequency greater than or equal to 2-5 Hz for greater than or equal to 5s (Twele et al., 2016a, 2017; Zeidler et al., 2018). The number and frequency of HVSWs per mouse were determined by developing an analysis pipeline, which included a step with a well-validated seizure detection algorithm obtained from the Krook-Magnuson lab (Zeidler et al., 2018).

Three hours of randomly selected data from the week prior to the FUS proof-of-concept study were chosen and passed through the SLE detection algorithm to confirm a minimum of 10 SLEs/hr for each mouse. During this review, electrographic behavioral seizures were also scored in the mice (n=2) using 72 randomized hours per mouse using the Racine scale, scoring any seizures observed using our in-house vEEG review software (Racine, 1975). Following FUS administration, the vEEG files were analyzed using our analysis pipeline, which included the seizure detection algorithm, to determine the effects of FUS stimulation on metrics of SLEs. Metrics evaluated were interevent duration of SLEs, SLE duration, and spike frequency within the SLE.

### Data analysis

Data from six mice from each group was analyzed to investigate if IHK and control mice had differences in behavioral training across the three stages in the number of sessions and days. An unpaired t-test was performed in Prism (Prism 10 for macOS, GraphPad) for each behavioral stage, comparing the two groups of mice for either number of sessions or days. The total number of sessions and days was also analyzed.

To evaluate the impact of FUS stimulation on SLE activity, the two mice from the proof-of-concept study were used as an example of the type of analysis that could be run to compare FUS to the Pre-FUS and Post-FUS session conditions for both stimulation target conditions and both FUS parameter sets. Using in-house MATLAB code, the metrics of SLEs were provided for each SLE identified per mouse per target condition and FUS parameter set run. GraphPad Prism was used to display the metrics of SLEs (interevent duration, SLE duration, and spike frequency) for the low FUS pressure and contralateral target condition for mouse 1.

### Histology

A subset of mice underwent transcardial perfusion using 4% paraformaldehyde (Thomas Scientific) in 1x phosphate buffer saline (Dulbecco’s). Afterward, the brains were kept in 4% paraformaldehyde for 24 hours, transferred to 30% sucrose (Sigma-Aldrich) for 48 hours, and then sectioned on a microtome (SM 2010 R, Leica) at a 60-micron thickness. The slices were then stained using a Cresyl Violet stain to distinguish the hippocampus for analysis (Umpierre et al., 2014). Hippocampal brain sections were viewed under brightfield microscopy (XL Core, EVOS) at 4x and viewed at the level of either the KA or saline injection coordinates to confirm pathophysiology for IHK mice.

## Results

### 3D printing provided a customizable and low-cost solution for head-fixation of mice

We developed a low-cost method to head-fix awake mice and administer FUS. This entailed creating a 3D-printed treadmill with head-fixing components. The head-fixing components included the headplates, a head-fixing arm, and pins (Figures 3 and 4A). Each headplate was designed to slide into the head-fixing arm(s) with a hole through which a pin could be inserted to prevent the mouse from releasing its head from the head-fixing arm. The treadmill housing apparatus consisted of a training platform, four legs, and a base (Figures 3B and 4B). The training platform had four tabs that were inserted into slots on the legs and included a cutout to make room for the foam wheel (White PE Foam Roller 8”x12” model 30-2261, CanDo) used as the surface for the mouse to run on. The training platform helped with behavioral training by providing an area for the mouse to adapt to walking around on the treadmill. The base consisted of four slots in the corners for the legs to insert into and provide stability. The components for both the treadmill and head-fixing are shown in Figures 3 and 4. The utilization of these custom parts led to a more efficient execution of the setup and provided greater flexibility for FUS administration. For example, the heights of the legs of the treadmill were adjusted to allow more room for positioning ultrasound equipment. Additionally, 3D printing components allowed for a low-cost method of head-fixing the mice compared to commercial products designed for the same purpose.

**Figure 4:**
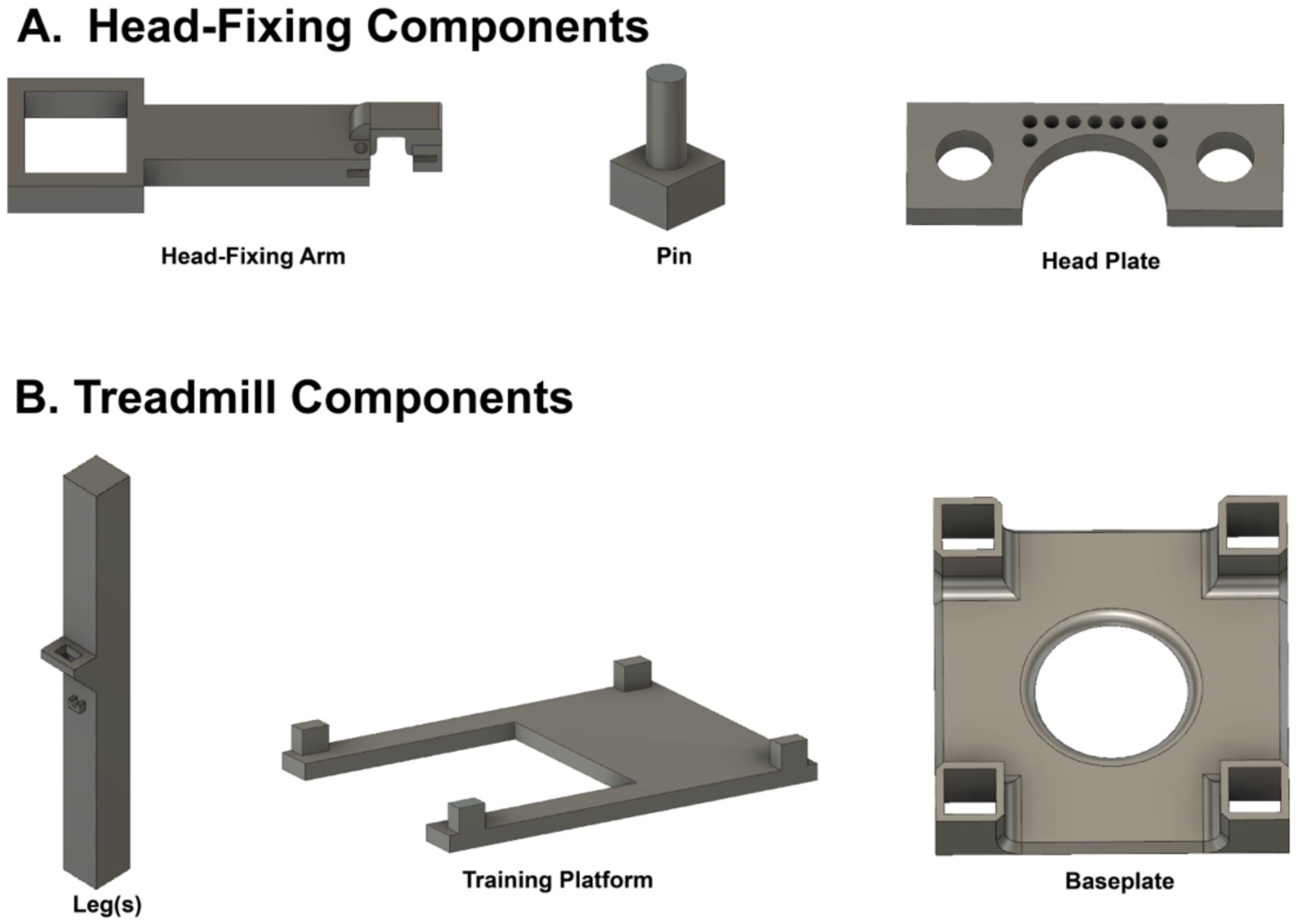
Design of custom-made components for head-fixing and restraining awake mice A) The image depicts the components from design files for the custom-made head-fixing components. The photo on the left shows the head fixing arm used in Figure 3 which was designed to allow the ultrasound transducer to come in on the side of the mouse’s head. The middle photo depicts the pin used to slide through the hole(s) of the head-fixing arm and head plate. The headplate is shown on the right. Holes were located at the top of the arc and allowed for dental cement to fill in the holes when attaching the plate to the mouse. This provided a more secure mount to the head. B) The image depicts the design for the custom-made treadmill components. The baseplate (right) had four rectangular slots in the corners for the legs to slide into for stability.

### IHK treated mice demonstrate hippocampal sclerosis and granule cell dispersion

Histological analysis was used to verify pathophysiology of the IHK model. An IHK mouse with verified HVSWs and behavioral seizures (n=1; Figure 5C and D) was compared to a saline control mouse (n=1; Figure 5A and B). Transections of the hippocampus were viewed under brightfield microscopy at 4x. IHK pathophysiology was confirmed as having granule cell layer dispersion, hilar interneuron and pyramidal cell loss, and hippocampal sclerosis (Figure 5C and D) (Maroso et al., 2011). Additionally, EEGs were reviewed and IHK mice (n=2) were shown to have an average of 2.33 behavioral seizures/day with an average Racine scale score of 3.66 across 72 randomized hours a week prior to the proof-of-concept study (Racine, 1975).

**Figure 5:**
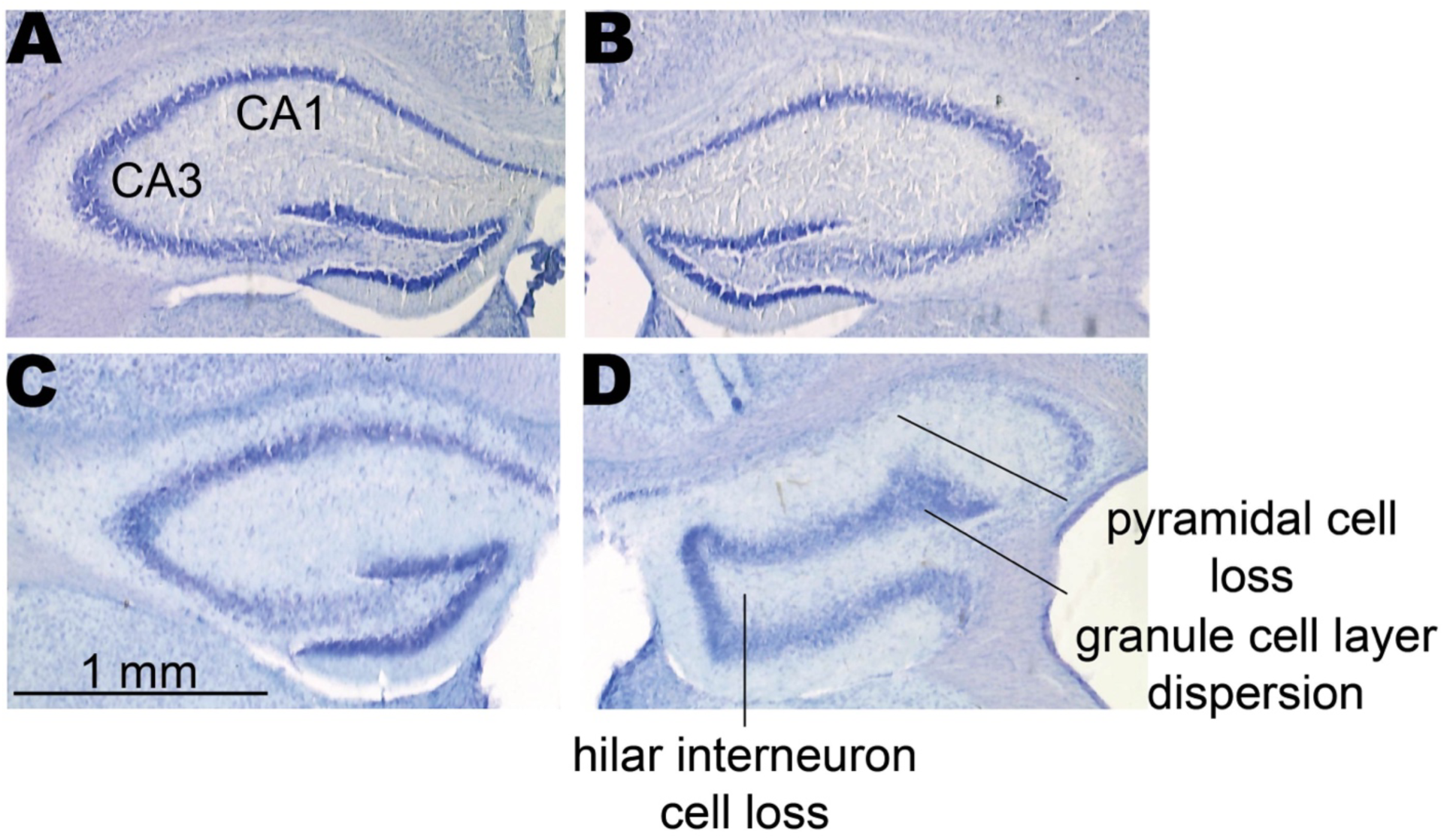
IHK administration causes damage in the hippocampus. A, B, show cresyl violet stained brain section obtained from a vehicle treated mouse, with the left and right hippocampus shown, respectively. The hippocampal CA1 and CA3 areas are labelled. C, D, show cresyl violet stained brain sections obtained from the contralateral and ipsilateral hippocampus from an IHK-treated mouse. Note the granule cell layer dispersion, pyramidal cell loss, hilar interneuron cell loss, and hippocampal sclerosis in the marked areas of CA1 and CA3 of the right hippocampus. This pathophysiology is representative of the IHK mouse (Maroso et al., 2011).

### IHK mice require more training sessions in a three-stage approach for the head-fixation apparatus

We developed three stages of behavioral training to acclimatize and head-fix the mice to the 3D-printed treadmill for FUS stimulation based off of Guo et al., 2014 (Guo et al., 2014). A sucrose solution was given in increments of 15-30 s to incentivize the mouse. The first stage was hand-holding (Figure 6A). After the mouse was accustomed to the handler’s hand, the training progressed to the next (second) stage: the platform (Figure 6B). The duration of the training sessions started with five minutes and increased to 10 minutes for both stages 1 and 2. The platform stage included familiarizing the mouse with the stage by placing it on the treadmill’s housing platform and foam wheel during the session. Once the mouse was comfortable with the stage and the foam wheel, the handler moved the mouse to the third training stage, the head-fixation stage (Figure 6C). In this phase of training, the handler attached the mouse’s headplate to the fixture and allowed the mouse to walk on the treadmill. When the mouse was adapted to head-fixation for 5 minutes, the handler increased the session time by increments of 10 and then 20 minutes, and so on until reaching 60 minutes. During this time, the mouse would run or walk at their own pace on the treadmill. While increasing the session time, the handler would wean the mouse off the sucrose reward by increasing the duration between rewards in increments of 30 s. The reward schedule was adapted from Guo et al. (Guo et al., 2014). Calm and comfortable behavior was characterized by mastery of the behavioral stage without fear, stress, or struggling. Training was considered complete when the mouse was comfortable on the treadmill for at least 60 minutes and could run or walk without encouragement from the handler. These three stages were used to train both IHK and control mice.

**Figure 6:**
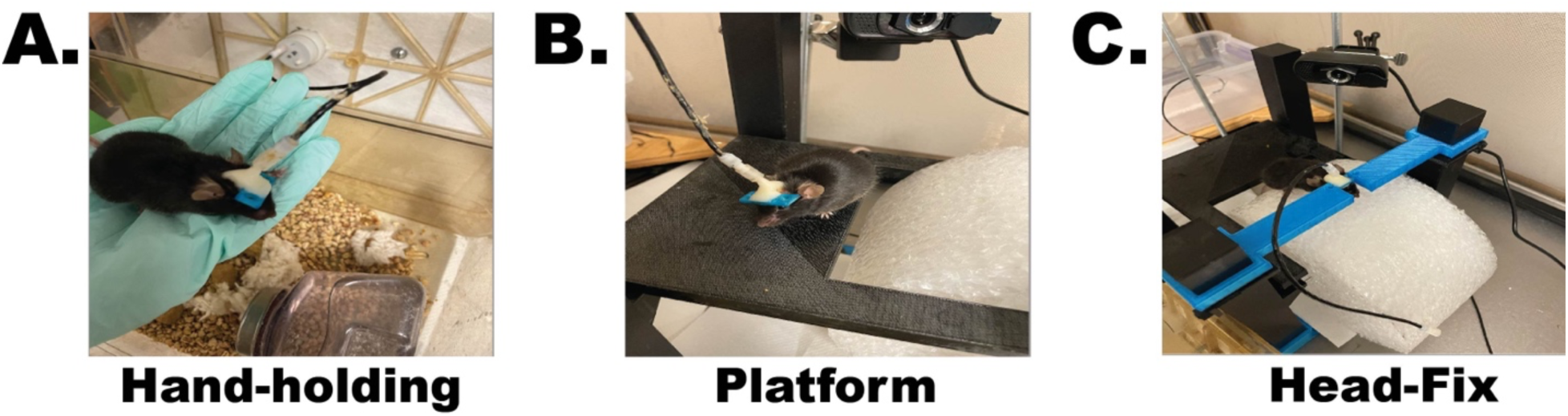
Behavioral training stages. The three stages of behavioral training are shown in A, B, and C as Hand-holding, Platform, and Head-Fix, respectively. The mouse would start with 5-minute sessions at each stage and 4-5 sessions a day until comfortable prior to moving to the next stage. When the mouse was adapted to head-fixation for 5 minutes, the handler increased the session time by increments of 10 and then 20 minutes, and so on until reaching 60 minutes. Training is considered completed when the mouse is comfortable on the treadmill and runs or walks without encouragement from the handler.

We compared IHK (n = 6) to control (n = 6) mice for each of the three behavioral stages for the number of sessions and days (Figure 7). The IHK mice showed an increase in the number of sessions needed for behavioral training compared to the control mice. The number of sessions required (Figure 7 Left) was significantly greater for the platform (p<0.05) and head-fix (p<0.01) stages, as well as the total number of sessions (p<0.001). While IHK mice took a greater number of sessions to train in all other stages of training, interestingly, in the hand-holding stage IHK mice took fewer sessions than the control mice to train. The number of days was significant (p<0.05) for the IHK mice when compared to the control mice for the platform stage only, with the IHK mice requiring more days to train. However, when looking at the total number of days, both groups of mice took the same number of days to complete the training, even though there was a difference in the total number of sessions to train. Overall, these findings suggest that there may be an impairment in learning in the IHK mice.

**Figure 7:**
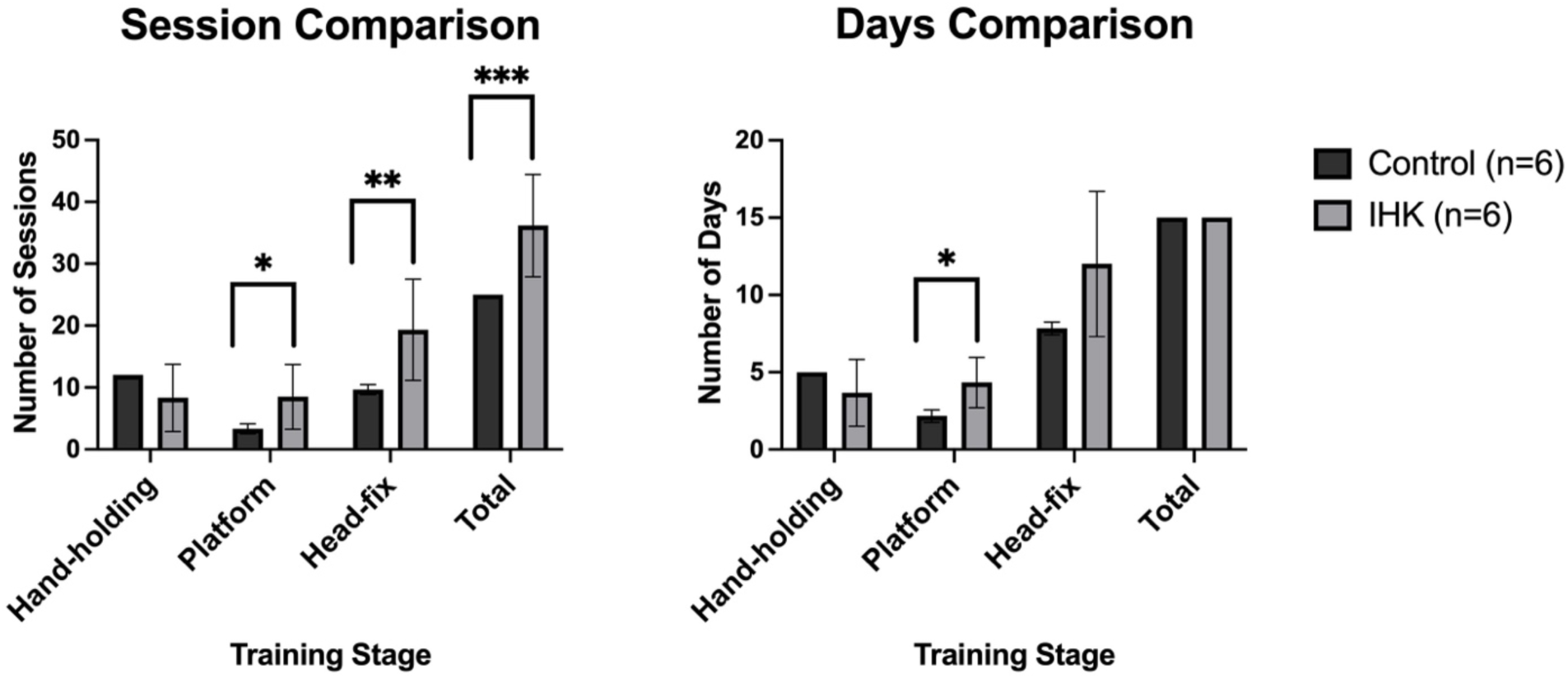
Behavioral training comparison between IHK (n=6) and control (n=6) mice by the number of sessions and days. Total number of sessions and days for the three stages of behavioral training, as well as total sessions and days are shown on the left and right, respectively. The IHK mice require more sessions to train, while the number of days required to successfully train remain the same when compared to control mice. Additionally, the IHK mice, when compared to the control mice, showed impairment in the Platform and Head-Fix stages in the number of sessions.

### Analysis pipeline allowed for visualization and verification of SLEs and extraction of metrics of SLEs recorded on EEG during FUS administration

An analysis pipeline was developed to evaluate FUS stimulation on metrics of SLEs (interevent duration, SLE duration, and spike frequency). Our analysis method was a multi-step process. First, the vEEG files were converted from the AcqKnowledge software format to the MATLAB format using the Bioread library in Python (3.8.8). Second, these converted MATLAB files were converted into a MATLAB naming convention and extracted information needed for the seizure detection algorithm was provided with our in-house code.

Third, these converted files were then loaded into the seizure detection algorithm in MATLAB (Student 2022a and 2023a, Mathworks) for further analysis (Zeidler et al., 2018). In order to detect SLEs, an automated electrographic seizure detection algorithm created by the Krook-Magnuson lab was used (Zeidler et al., 2018). The seizure detection algorithm allowed users to adjust several settings and create a seizure classification that the seizure detection algorithm would search EEG data for and flag qualifying events. The seizure detection algorithm had a graphical user interface which allowed the user to adjust and input settings and parameters for their specific needs and analysis. We chose to adjust a threshold setting, seizure frequency and length settings, and additional noise and overlapping event settings. The baseline setting was set to the default values which meant that spikes in the EEG data had to exceed the amplitude of 90% of the data in the EEG file to be flagged by the seizure detection algorithm. The threshold setting was calculated and manually set to twice the baseline value (Duveau et al., 2016; Twele et al., 2016b). This meant that a spike in the EEG data had to meet twice the baseline value or exceed it. For seizure frequency and length values the classification criteria for HVSWs was used (Twele et al., 2016a; Zeidler et al., 2018). For the noise and overlapping event settings, the option to capture both positive, negative, and overlapping events was selected, and a “split noise” option was checked that would prevent events containing noise from being counted as an SLE. A “Glue” events setting which connected events with a predetermined interevent interval was set to three seconds (Twele et al., 2016a). Once the seizure detection algorithm identified SLEs, “peak width” was adjusted per mouse mostly between 0.5 – 0.1 seconds to eliminate spikes/peaks that were being included but were noise. The data was rerun through the seizure detection algorithm to ensure a “clean” report of finalized detected SLEs. The seizure detection algorithm facilitated the identification of SLEs in a list format based on our criteria alone but did not allow for the visualization and verification of SLEs. Visualization of the EEG data along with the duration of the detected SLE and spike frequency of the SLE allows for verification against set criteria and field standard (Twele et al., 2016a, 2016b).

Fourth and finally, to allow for visualization and verification of SLEs, custom in-house review software was developed in MATLAB that allowed for the identified SLEs along with their EEG component, duration of the SLE, and spike frequency of the SLE to be reviewed by a trained reviewer and verified as SLEs. The trained reviewer can choose a predetermined baseline period in seconds before and after the SLE to aid in their visualization. During this review process, metrics of SLEs were then determined and recorded for each verified SLE. As there were many custom-built add-ons to the SLE detection algorithm, a false positive/negative rate was conducted to validate the analysis pipeline. Evaluating 10 randomized experimental sessions for two IHK mice in the pilot study for a total of 242 SLEs, the false positive and negative rates were 1% and 7.86% respectively. Detected seizures from the detection algorithm were compared against the raw EEG files by an expert reviewer. We created a multi-step approach to our analysis pipeline for detecting and verifying SLEs and extracting metrics of SLEs. Our custom-made code is available at https://github.com/wilcox-lab/FUS_Neuromodulation_Code.

### SLE metrics can be evaluated during FUS sessions

In order to test the ability of SLEs to be recorded during FUS administration, trained mice were head-fixed, and baseline SLEs were recorded for 60 minutes before delivering low pressure FUS to the contralateral hippocampus for 40 minutes. This was followed by recording EEGs for an additional 60 minutes. SLEs were then reviewed before, during, and after FUS stimulation (Figure 8). Identified SLEs using our in-house visualization code are indicated in red and are distinguished by the high spiking frequency and high amplitude from baseline EEG levels. Our visualization code allows us to see SLE duration, number of spikes in the SLE, and spike frequency of the detected SLE to help the expert reviewer in classifying the detected SLE. Once the SLE is verified (i.e. criteria checked as HVSW and EEG signal checked) the metrics of the SLE are saved for analysis. The effect of FUS on all metrics of SLEs (interevent duration, SLE duration, and spike frequency) is shown in Figure 9 and discussed below.

**Figure 8:**
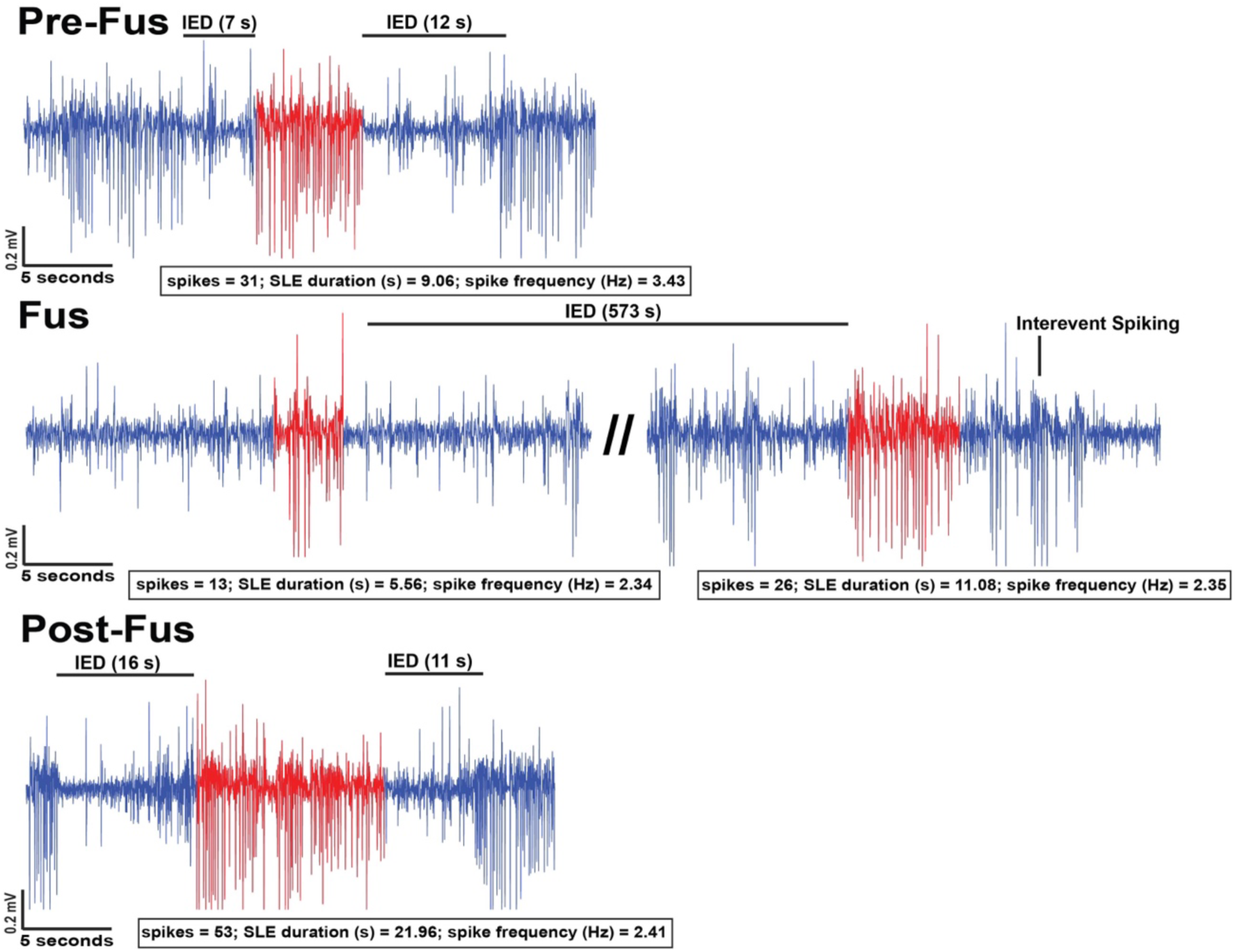
Representative SLE traces during FUS administration. From top to bottom, the EEG traces are organized by session condition, from Pre-FUS, FUS, and Post-FUS. The low pressure FUS parameter set and contralateral target condition were used in this example. In red are examples of SLEs identified that meet the criteria for a HVSW. Number of spikes, SLE duration (s), and spike frequency (Hz) of each SLE highlighted in red are displayed below each EEG trace. Interevent duration (IED) is marked on the figure, and shows that there is an increase of IED during FUS. Interevent spiking is also noted in the figure, however not analyzed.

**Figure 9:**
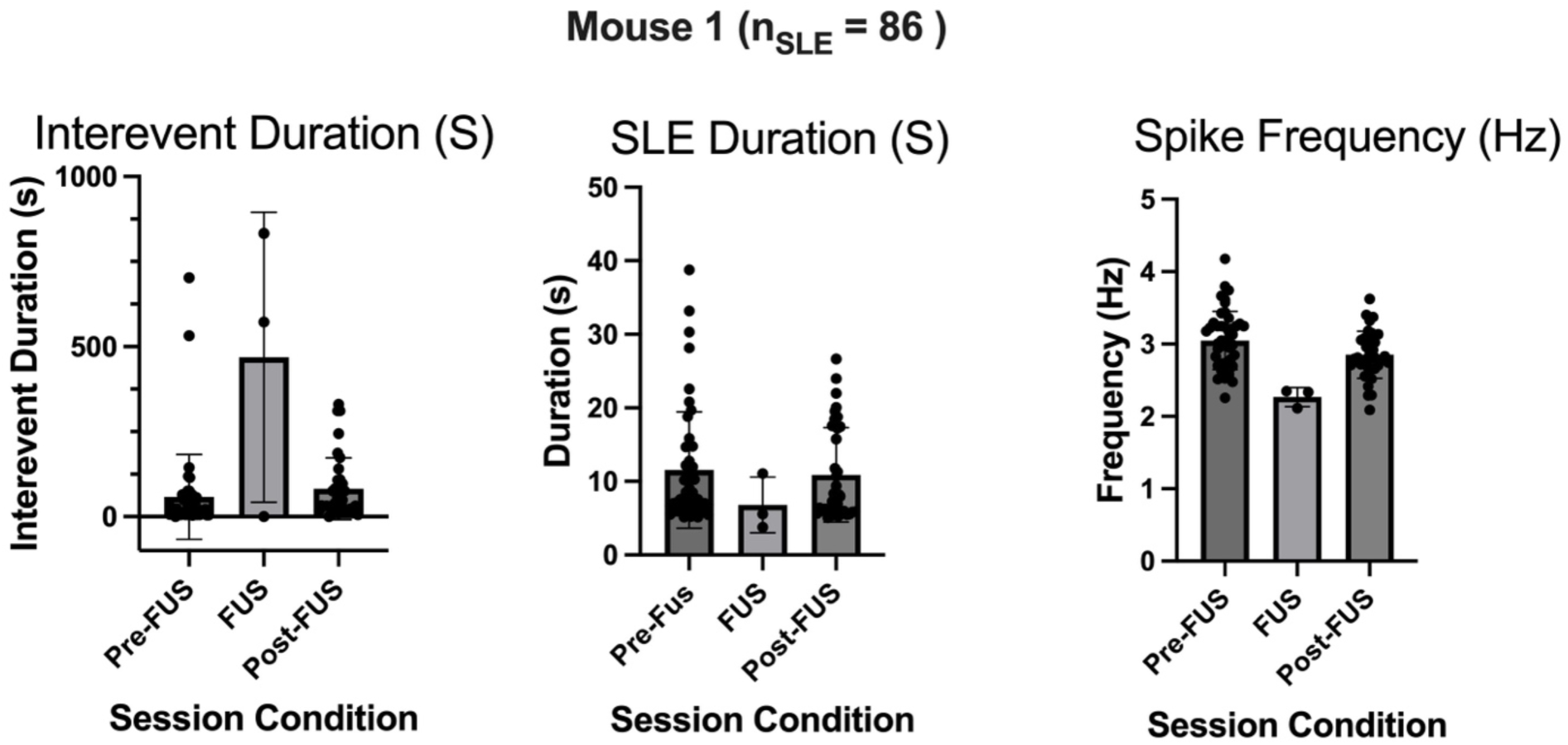
Representative analysis for metrics of SLEs from FUS stimulation. Displayed in this figure are the metrics of SLEs (interevent duration, SLE duration, and spike frequency) for the low pressure FUS parameter set and contralateral target condition for mouse 1. Interevent duration, SLE duration, and spike frequency for each SLE were determined for each session condition (Pre-FUS, FUS, Post-FUS) (n=86 SLEs).

An example of the data using our analysis method is displayed in Figure 9. Here, we show the metrics of SLEs for mouse 1, with each SLE being a data point. A total of 86 SLEs were detected in the example for the low pressure FUS parameter set and contralateral target condition. In this condition, FUS increases interevent duration and decreases spike frequency during FUS stimulation. FUS has a noticeable difference on the time between SLEs during stimulation and even carries an effect Post-FUS. Figure 8 gave an example set of SLEs being compared. We can see the increase in interevent duration from Pre-FUS to FUS was 12 s to 573 s, with a return to 11 s at Post-FUS. Whereas, in Figure 9, the averages of all SLEs were taken. The increase in interevent duration from Pre-FUS to FUS was from 58 s to 469 s, with a return to 82 s at Post-FUS. SLE duration decreased from 12 s during Pre-FUS to 7 s during FUS and returning to 11 s during Post-FUS. In this representative example, spike frequency decreased from 3 Hz in Pre-FUS to 2.3 Hz during FUS returning to 2.9 Hz in the Post-FUS condition. While FUS may show some promise in decreasing spiking activity in this example, the effects are modest and may not be biologically relevant. However, future work may determine if FUS can effectively modulate seizure activity in a larger population of mice.

We have provided all the metrics of SLEs across all FUS parameter sets and stimulation target conditions for both mice in the proof-of-concept study in Figures S1 – S3. Interevent duration, SLE duration, and spike frequency are shown in S1, S2, and S3 respectively. Data from the low pressure FUS parameter set with bilateral targeting (Low Bilateral) is shown top left. The low pressure FUS parameter set with contralateral targeting (Low Contralateral) is shown in the top right panel. The high pressure FUS parameter set with bilateral targeting (High Bilateral) is shown in the bottom left panel. The high pressure FUS parameter set, with contralateral targeting (High Contralateral) is shown in the bottom right panel. Looking at all metrics, the interevent duration shows the most interesting findings in the contralateral stimulation target condition, regardless of low or high pressure, where we see that during FUS stimulation, interevent duration increases, suggesting a reduction of SLE incidence. However, future studies in a larger cohort will be needed to conclude that FUS stimulation can reduce seizure incidence in IHK mice.

### A method to administer FUS to awake, head-fixed mice with TLE while simultaneously recording EEG

We developed a method to administer FUS to awake, head-fixed mice with TLE while simultaneously recording vEEG. Our innovative approach is summarized in Figure 10, with a block diagram of the setup and an example pulse waveform of the ultrasonic stimulation. We used MATLAB to program the function generator to produce the desired ultrasonic waveforms, which were sent through an RF amplifier and then to the ultrasound transducer to deliver FUS to the mouse. Additionally, a copy of the ultrasound waveform signal was sent from the function generator to another amplifier and recorded back through the EEG acquisition software package to ensure accurate timing of stimulation pulse with the EEG recording from the implanted electrode from the mice. The mice have been trained to walk/run on the 3D printed low-cost treadmill during the FUS administration. We have shown that mice can tolerate our setup.

**Figure 10:**
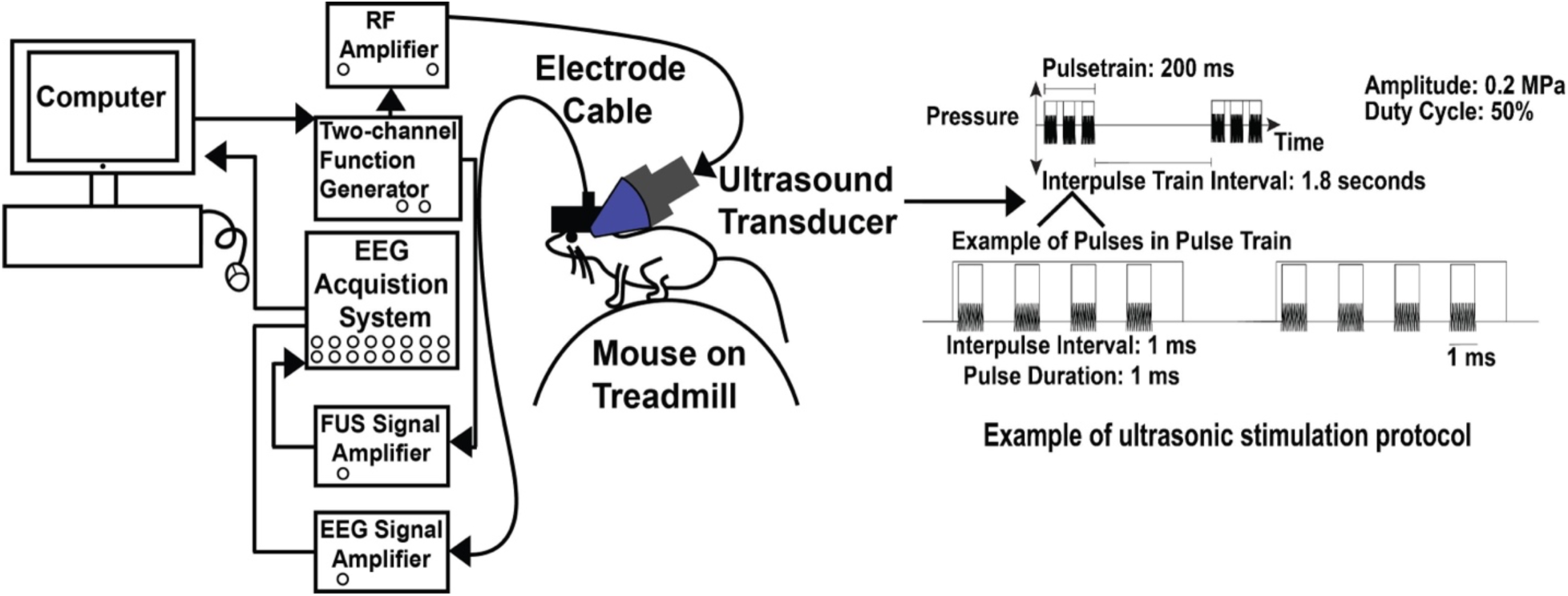
Block diagram of our method to deliver FUS in awake, head-fixed mice with simultaneous EEG. This figure shows a block diagram of the method workflow, starting with the computer sending out a programmed waveform to the two-channel function generator. The function generator outputs through the RF amplifier to the ultrasound transducer in order to stimulate the mouse. The image depicts a mouse undergoing bilateral target stimulation condition while positioned on a custom treadmill. On the left side of the mouse’s head, there is a FUS collimator printed in clear resin, which is attached to the FUS transducer. Additionally, the mouse is head-fixed to a 3D-printed treadmill, which is depicted in the block diagram as a cartoon. Through the second channel the ultrasound stimulation waveform was sent to a FUS amplifier for EEG acquisition. The mouse’s EEG signal was also recorded. The example ultrasound stimulation protocol is of the low pressure FUS parameter set of 0.2 MPa. Shown here is the pulse train duration of 200 ms with the interpulse train interval of 1.8 s. Additionally, this FUS parameter set had a 50% duty cycle and used the 500 kHz transducer.

## Discussion

FUS has exciting potential for the treatment of people with drug-resistant epilepsy, as it is non-invasive and provides extraordinary spatial resolution and target specificity for neuromodulation (Mahoney et al., 2020). However, safety and efficacy should be determined in preclinical models of epilepsy before attempting this approach in humans. Previous work in the field was largely limited by the use of anesthetized animals that did not have epilepsy but had evoked seizures. Thus, to advance the field, the work described here provides a cost-effective method to study FUS in awake, head-fixed mice with TLE, while simultaneously recording vEEG.

We used our approach to study the effects of behavioral training in IHK and control mice across three behavioral training stages, wherein we determined the number of sessions and days required for training. IHK mice were significantly impaired in behavioral training in the platform and head-fixed stages in the number of sessions and total sessions. However, the total number of training days remained constant between the two groups of mice. This suggests that the IHK mice were either cognitively impaired or, perhaps, had more anxiety than the control mice. Both cognitive impairment and anxiety are often observed in rodent models of epilepsy (Zeidler et al., 2018).

Next, we used our setup to provide a proof-of-principle data set. We determined the metrics of SLEs for the low pressure FUS parameter set and contralateral targeting. In this example, FUS had the most effect on interevent duration by increasing the time between events. However, further testing on a larger cohort is needed to confirm this effect.

FUS is known to have limitations when administered in rodents. Studying FUS in rodents was difficult to control for the auditory confound. The auditory confound/artifact is when FUS stimulates the auditory network, and without a proper control it may be difficult to discern if FUS produces a direct neuromodulation effect or auditory effect (Cornelssen et al., 2023). There have been numerous approaches to control for this confound, such as a no-stimulation control or applying an envelope to the waveform (Cornelssen et al., 2023). More recently, a transgenic mouse line was developed that provides a “clean” mouse for ultrasound (Guo et al., 2023). In this mouse line, diphtheria toxin was used to destroy hair cells in the ears. This renders the mouse deaf and is thus a way to control for any auditory artifact that might occur as a consequence of FUS delivery (Guo et al., 2023). Additionally, in rodents, there is the potential of stimulating more brain area than the intended target. In rodents, the pressure field can spread out over the small brain area. One way to overcome this limitation is to move away from rodent models to a larger animal model, such as a guinea pig model of epilepsy.

Another limitation of FUS administration in rodents while simultaneously recording EEG can be that the ultrasound stimulation could cause artifacts in EEG recording due to direct electric coupling or auditory, tactile, or vestibular artifacts. In this study, we aimed to minimize these issues by coupling the ultrasound transducer to the opposite side of the head from the electrode implant. Additionally, we limited the I_SPTA_ to at or below 333.33 mW/cm^2^. Nonetheless, only effects following the stimulation should be considered artifact-free. The effects reported during the stimulation might be affected by an artifact despite the above controls.

We have successfully developed an approach to deliver FUS to awake, head-fixed mice with TLE while simultaneously recording vEEG. We have shown in the proof-of-concept study that the mice can tolerate the setup and the data can be analyzed using our analysis pipeline. This optimized and cost-effective approach sets the stage for future preclinical experiments investigating the ability of FUS to decrease seizures through non-invasive neuromodulation.

## Acknowledgments

The authors would like to thank Drs. Daria Anderson, Kyle Thomson, Taylor Webb, and Peter West, as well as Jerry Saunders and Matthew Wilson for their feedback and helpful discussions on this project. We would like to thank Drs. Carlos Rueda and Jose Reyes for their assistance with mouse surgeries. Additionally, the authors would like to thank Drs. Esther Krook-Magnuson and Martha Streng on their assistance with the seizure detection algorithm. Finally, the authors would like to thank Drs. Amy Brooks-Kayal and Esther Krook-Magnuson for helpful discussions while implementing the IHK model. This work has appeared in the dissertation of Carena Cornelssen (Cornelssen, 2024).

## Data Availability Statement

The datasets and design files for this study are available upon request. Our analysis code is available at https://github.com/wilcox-lab/FUS_Neuromodulation_Code.

## Author Contributions

CC, KW, and JK conceived the project. CC, BB, and EF developed the methodology. CC, BB, EF, SB, and SS collected data. CC and EF analyzed data. CC, BB, and SS wrote the manuscript. CC, BB, EF, and KSW revised the manuscript. All authors approved the final submission.

## References

Chen, S. G., Tsai, C. H., Lin, C. J., Lee, C. C., Yu, H. Y., Hsieh, T. H., et al. (2020). Transcranial focused ultrasound pulsation suppresses pentylenetetrazol induced epilepsy in vivo. Brain Stimul 13, 35–46. doi: 10.1016/j.brs.2019.09.011

Constans, C., Deffieux, T., Pouget, P., Tanter, M., and Aubry, J. F. (2017). A 200-1380-kHz Quadrifrequency Focused Ultrasound Transducer for Neurostimulation in Rodents and Primates: Transcranial in Vitro Calibration and Numerical Study of the Influence of Skull Cavity. IEEE Trans Ultrason Ferroelectr Freq Control 64, 717–724. doi: 10.1109/TUFFC.2017.2651648

Cornelssen, C. (2024). Development of focused ultrasound (FUS) approaches to modify neural circuits underlying seizures in temporal lobe epilepsy (TLE). Salt Lake City: University of Utah.

Cornelssen, C., Finlinson, E., Rolston, J. D., and Wilcox, K. S. (2023). Ultrasonic therapies for seizures and drug-resistant epilepsy. Front Neurol 14, 1301956. doi: 10.3389/FNEUR.2023.1301956/BIBTEX

Duveau, V., Pouyatos, B., Bressand, K., Bouyssières, C., Chabrol, T., Roche, Y., et al. (2016). Differential Effects of Antiepileptic Drugs on Focal Seizures in the Intrahippocampal Kainate Mouse Model of Mesial Temporal Lobe Epilepsy. CNS Neurosci Ther 22, 497–506. doi: 10.1111/cns.12523

Engel, J., McDermott, M. P., Wiebe, S., Langfitt, J. T., Stern, J. M., Dewar, S., et al. (2012). Early surgical therapy for drug-resistant temporal lobe epilepsy: A randomized trial. JAMA - Journal of the American Medical Association 307, 922–930. doi: 10.1001/jama.2012.220

Guo, H., Salahshoor, H., Wu, D., Yoo, S., Sato, T., Tsao, D. Y., et al. (2023). Effects of focused ultrasound in a “clean” mouse model of ultrasonic neuromodulation. iScience 26, 108372. doi: 10.1016/J.ISCI.2023.108372

Guo, Z. V., Hires, S. A., Li, N., O’Connor, D. H., Komiyama, T., Ophir, E., et al. (2014). Procedures for Behavioral Experiments in Head-Fixed Mice. PLoS One 9, e88678. doi: 10.1371/journal.pone.0088678

Hakimova, H., Kim, S., Chu, K., Lee, S. K., Jeong, B., and Jeon, D. (2015). Ultrasound stimulation inhibits recurrent seizures and improves behavioral outcome in an experimental model of mesial temporal lobe epilepsy. Epilepsy and Behavior 49, 26–32. doi: 10.1016/j.yebeh.2015.04.008

Hesdorffer, D. C., Logroscino, G., Benn, E. K. T., Katri, N., Cascino, G., and Hauser, W. A. (2011). Estimating risk for developing epilepsy: A population-based study in Rochester, Minnesota. Neurology 76, 23–27. doi: 10.1212/WNL.0b013e318204a36a

Li, X., Yang, H., Yan, J., Wang, X., Li, X., and Yuan, Y. (2019a). Low-Intensity Pulsed Ultrasound Stimulation Modulates the Nonlinear Dynamics of Local Field Potentials in Temporal Lobe Epilepsy. Front Neurosci 13, 287. doi: 10.3389/fnins.2019.00287

Li, X., Yang, H., Yan, J., Wang, X., Yuan, Y., and Li, X. (2019b). Seizure control by low-intensity ultrasound in mice with temporal lobe epilepsy. Epilepsy Res 154, 1–7. doi: 10.1016/j.eplepsyres.2019.04.002

Lin, Z., Meng, L., Zou, J., Zhou, W., Huang, X., Xue, S., et al. (2020). Non-invasive ultrasonic neuromodulation of neuronal excitability for treatment of epilepsy. Theranostics 10, 5514–5526. doi: 10.7150/thno.40520

Mahoney, J. J., Hanlon, C. A., Marshalek, P. J., Rezai, A. R., and Krinke, L. (2020). Transcranial magnetic stimulation, deep brain stimulation, and other forms of neuromodulation for substance use disorders: Review of modalities and implications for treatment. J Neurol Sci 418, 117149. doi: 10.1016/J.JNS.2020.117149

Maroso, M., Balosso, S., Ravizza, T., Iori, V., Wright, C. I., French, J., et al. (2011). Interleukin-1β Biosynthesis Inhibition Reduces Acute Seizures and Drug Resistant Chronic Epileptic Activity in Mice. Neurotherapeutics 8, 304. doi: 10.1007/S13311-011-0039-Z

Min, B. K., Bystritsky, A., Jung, K. I., Fischer, K., Zhang, Y., Maeng, L. S., et al. (2011). Focused ultrasound-mediated suppression of chemically-induced acute epileptic EEG activity. BMC Neurosci 12, 23. doi: 10.1186/1471-2202-12-23

Murphy, K. R., Farrell, J. S., Gomez, J. L., Stedman, Q. G., Li, N., Leung, S. A., et al. (2022). A tool for monitoring cell type-specific focused ultrasound neuromodulation and control of chronic epilepsy. doi: 10.1073/pnas

Paxinos, G., and Franklin, K. B. J. (2001). The mouse brain in stereotaxic coordinates. San Diego.

Racine, R. J. (1975). Modification of seizure activity by electrical stimulation: Cortical areas. Electroencephalogr Clin Neurophysiol 38, 1–12. doi: 10.1016/0013-4694(75)90204-7

Thomson, K. E., and White, H. S. (2014). A novel open-source drug-delivery system that allows for first-of-kind simulation of nonadherence to pharmacological interventions in animal disease models. J Neurosci Methods 238, 105–111. doi: 10.1016/J.JNEUMETH.2014.09.019

Twele, F., Schidlitzki, A., Töllner, K., and Löscher, W. (2017). The intrahippocampal kainate mouse model of mesial temporal lobe epilepsy: Lack of electrographic seizure-like events in sham controls. Epilepsia Open 2, 180–187. doi: 10.1002/epi4.12044

Twele, F., Töllner, K., Bankstahl, M., and Löscher, W. (2016a). The effects of carbamazepine in the intrahippocampal kainate model of temporal lobe epilepsy depend on seizure definition and mouse strain. Epilepsia Open 1, 45–60. doi: 10.1002/epi4.2

Twele, F., Töllner, K., Brandt, C., and Löscher, W. (2016b). Significant effects of sex, strain, and anesthesia in the intrahippocampal kainate mouse model of mesial temporal lobe epilepsy. Epilepsy and Behavior 55, 47–56. doi: 10.1016/j.yebeh.2015.11.027

Umpierre, A. D., Remigio, G. J., Dahle, E. J., Bradford, K., Alex, A. B., Smith, M. D., et al. (2014). Impaired cognitive ability and anxiety-like behavior following acute seizures in the Theiler’s virus model of temporal lobe epilepsy. Neurobiol Dis 64, 98–106. doi: 10.1016/j.nbd.2013.12.015

Walton, D., Spencer, D. C., Nevitt, S. J., and Michael, B. D. (2021). Transcranial magnetic stimulation for the treatment of epilepsy. Cochrane Database Syst Rev 2021. doi: 10.1002/14651858.CD011025.PUB3

Zeidler, Z., Brandt-Fontaine, M., Leintz, C., Krook-Magnuson, C., Netoff, T., and Krook-Magnuson, E. (2018). Targeting the mouse ventral hippocampus in the intrahippocampal kainic acid model of temporal lobe epilepsy. eNeuro 5, 158–176. doi: 10.1523/ENEURO.0158-18.2018

Zou, J., Meng, L., Lin, Z., Qiao, Y., Tie, C., Wang, Y., et al. (2020). Ultrasound Neuromodulation Inhibits Seizures in Acute Epileptic Monkeys. iScience 23. doi: 10.1016/j.isci.2020.101066

