## Supplementary Figures for "A method for focused ultrasound (FUS) neuromodulation with simultaneous electroencephalogram recordings in awake, head-fixed mice with temporal lobe epilepsy": Supplementary_Material_3.19.24_biorxiv.pdf

### 1 Supplementary Figures and Tables

#### 1.1 Supplementary Figures

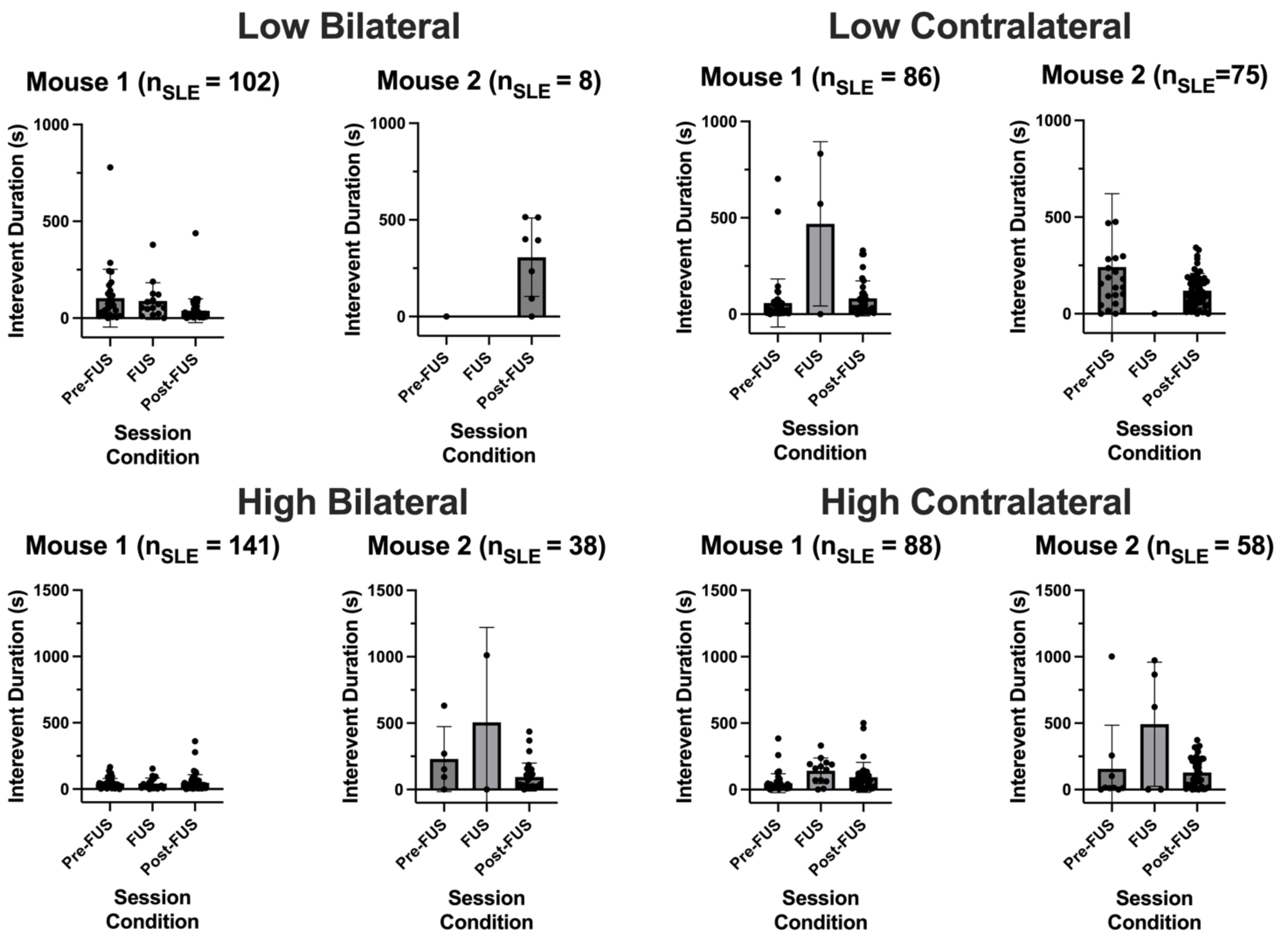


**Supplementary Figure 1.** Interevent Duration (S). The interevent duration (s) is displayed for both mouse 1 and 2 for each FUS parameter set (Low and High) and stimulation target (Bilateral and Contralateral) conditions. The interevent duration is displayed for each SLE for each session condition (Pre-FUS, FUS, Post-FUS). Note that the y-axis scale changes between conditions.

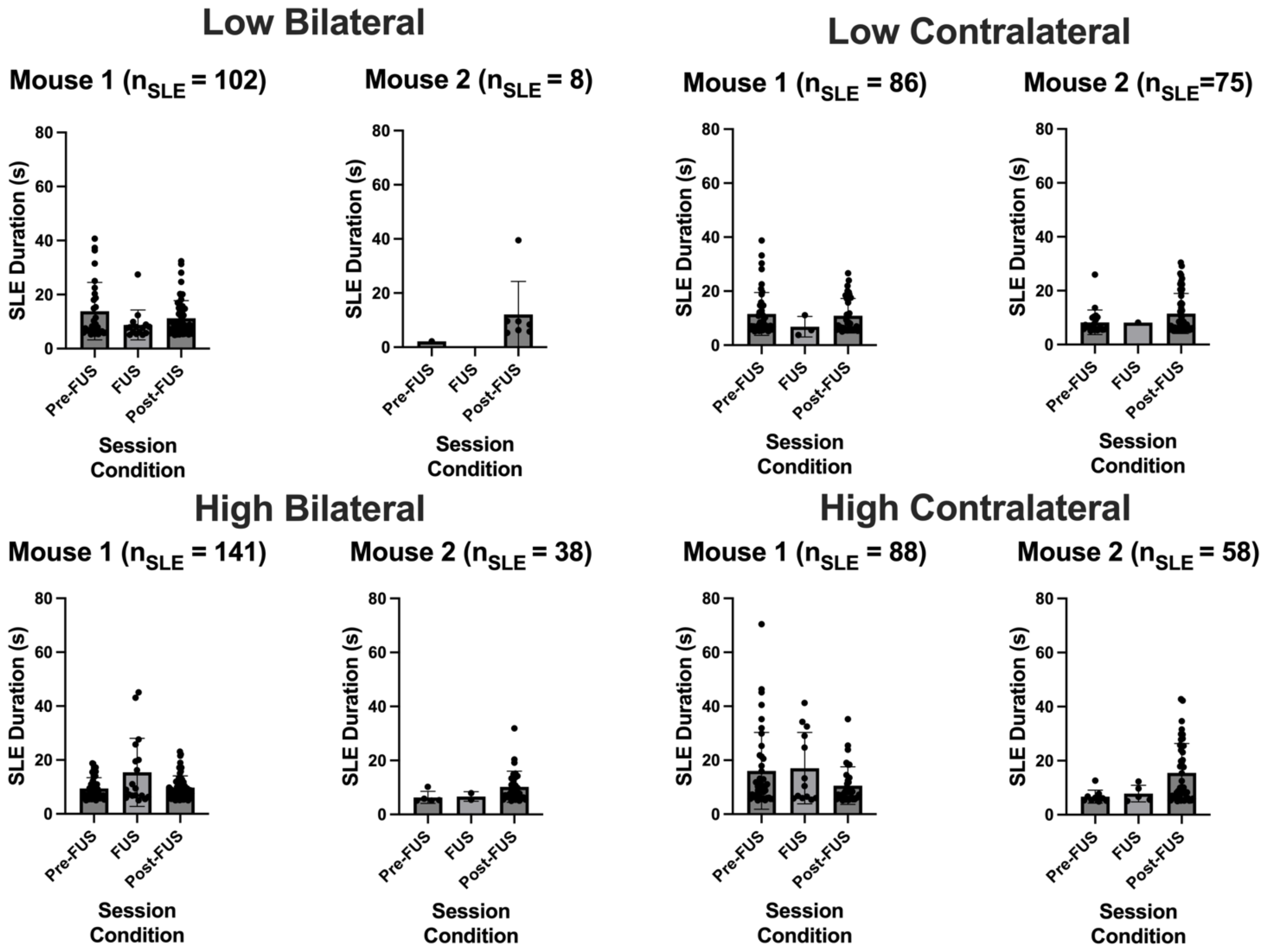


**Supplementary Figure 2.** SLE Duration (S). The SLE duration (s) is displayed for both mouse 1 and 2 for each FUS parameter set (Low and High) and stimulation target (Bilateral and Contralateral) conditions. The SLE duration is displayed for each SLE for each session condition (Pre-FUS, FUS, Post-FUS).

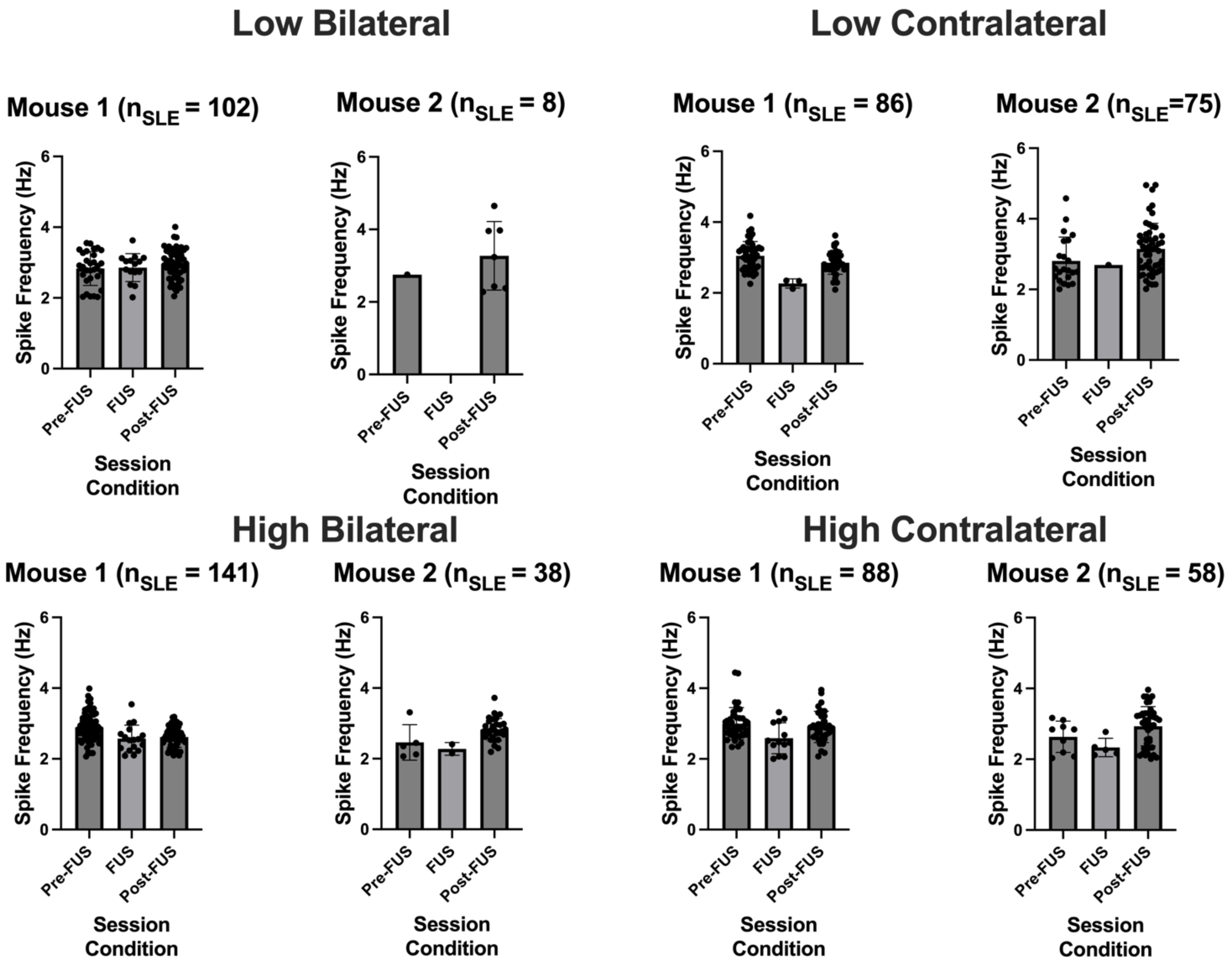


**Supplementary Figure 3.** Spike Frequency (Hz). The spike frequency (Hz) is displayed for both mouse 1 and 2 for each FUS parameter set (Low and High) and stimulation target (Bilateral and Contralateral) conditions. The spike frequency is displayed for each SLE for each session condition (Pre-FUS, FUS, Post-FUS).
